# Noninvasive detection of Phenylalanine in the human brain with MRS at 7T

**DOI:** 10.1101/2023.10.16.562609

**Authors:** Nouha Salibi, Thomas S Denney, Gopikrishna Deshpande

## Abstract

We set out to measure phenylalanine in the human brain using magnetic resonance spectroscopy (MRS) in a ultra-high field 7T MRI scanner. Phenylalanine is a precursor to Norepinephrine, a neurotransmitter important for attention and arousal. Depletion in norepinephrine, especially in the locus coeruleus, has been implicated as an etiological factor in Alzheimer’s disease. Therefore, being able to noninvasively measure phenylalanine *in vivo* in humans has a multitude of translational applications. Using phantom experiments, we first validate and optimize the MRS techniques used for observing phenylalanine in the brain at 7T. However, we failed to detect phenylalanine in human volunteers (N=15). In order to understand the reasons for this failure, we performed experiments in a cat model with external phenylalanine injections to determine the amount of phenylalanine required for it to be detected *in vivo* in the brain. This threshold was found to be 3.4 mM. This indicated that phenylalanine concentrations in both healthy and AD patients, that too in a small region such as the locus coeruleus, will likely not meet this threshold. Therefore, we conclude that even with state-of-the-art technologies and 7T MRI, it is not possible to detect phenylalanine in the human brain *in vivo* under natural conditions.

## INTRODUCTION

Proton MR spectroscopy (1H MRS) is a non-invasive technique for evaluation of *in vivo* metabolites that are not seen on MR (Magnetic Resonance) images, which are generated from strong water and fat signals. MRS is routinely used to investigate and monitor metabolic abnormalities related to brain pathology. The most commonly observed brain metabolites include NAA (N-Acetyl aspartate), choline (Cho), creatine (Cr), myo-inositol (mI) and elevated lactate. Metabolites with *in vivo* concentrations of 1 *mmol* or greater are routinely measured on 1.5T and 3T clinical scanners. Higher field strength scanners can detect lower concentration of metabolites. We speculated that it may be possible to detect *in vivo* phenylalanine at 7T and determine the reliability of its quantification. As far as we know, there has been no attempt at observing phenylalanine at 7T in normal or diseased brain at endogenous concentrations. All spectroscopy 7T studies were focused on better quantification of small concentration metabolites upfield of water such as NAA, Cr, Gho, glutamine, Glutamate, GABA, and myo-inositol (illustration in Fig.1).

**Fig 1:**
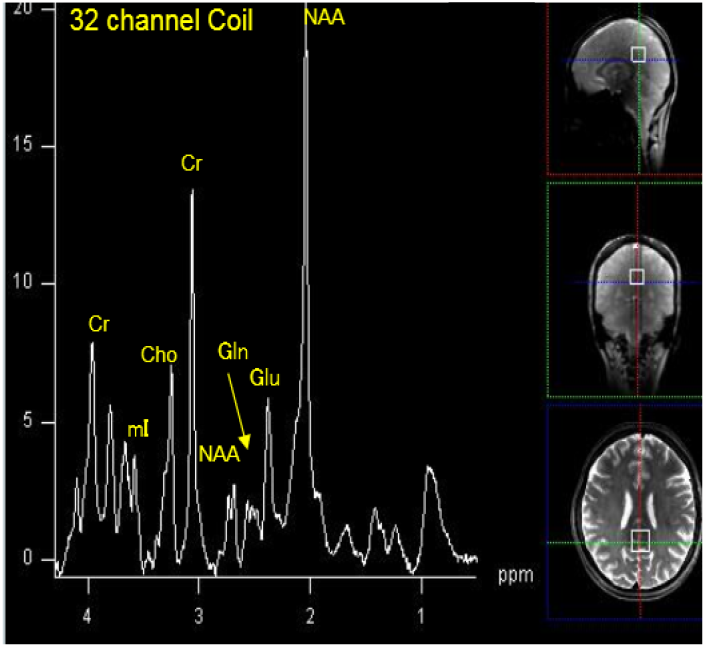
Typical Brain spectrum from a 8 cm3 voxel (white box) acquired with a 32 channel head coil at 7T. The spectrum displays standard metabolite peaks upfield from (to the right) of the water peak at 4.7 ppm (not displayed).

Phenylalanine is an amino-acid that is known under 3 forms: L-phenylalanine, which is found naturally in proteins; D-phenylalanine, which is a mirror image of L-phenylalanine made in the laboratory; and a DL-phenylalanine which is a combination of the L- and D-phenylalanine. *In vivo*, phenylalanine is transformed into the amino-acid tyrosine as well as metabolites such as L-dopa, epinephrine, norepinephrine. It is precisely this property of phenylalanine that makes it a valuable metabolite to measure since it is a precursor to norepinephrine, an important neurotransmitter in the brain. In healthy individuals, norepinephrine increases alertness, arousal, and attention, as well as enhances formation of memories. However, depletion of norepinephrine in the locus coeruleus in the brain stem has been shown to be an etiological factor in the development of mild cognitive impairment (MCI) and Alzheimer’s disease (AD).

Locus Coeruleus in the brain stem is the largest repository of norepinephrine in the human brain [1]. Noradrenergic neurons in the locus coeruleus project to many parts of the brain including olfactory, limbic and prefrontal areas. Norepinephrine is known to suppress neuroinflammation [2]. This purported role has been hypothesized to be a protective factor against AD. In fact, Heneka et al [3] showed that norepinephrine stimulation of mouse microglia suppressed Aβ-induced cytokine and chemokine production and increased microglial migration and phagocytosis of Aβ. Induced degeneration of LC increased expression of inflammatory mediators in APP-transgenic mice and resulted in elevated Aβ deposition. This indicates that decrease of norepinephrine in LC facilitates the inflammatory reaction of microglial cells in AD and impairs microglial migration and phagocytosis, thereby contributing to reduced Aβ clearance. These findings indicate that depletion of norepinephrine in LC is an etiological factor in the development of MCI and progression to AD.

L-phenylalanine structure C_6_H_5_CH_2_CH(NH_2_)COOH, is shown in Fig 2. Its high-resolution spectrum (Fig 2) consists of a multiplet between 7.3 and 7.45 ppm from the protons in the phenyl ring, and smaller resonances in the form of doublets of doublets at 3.11ppm, 3.28 ppm (from CH2) and 3.98 ppm (from CH). Measuring in vivo phenylalanine is challenging because of its very small concentration in the normal brain and because phenylalanine peaks overlap with other metabolites that have much higher concentrations. The aim of the current study is to measure the dominant phenylalanine resonance between 7.3 and 7.45 ppm for testing the feasibility and reliability of phenylalanine detection and quantification at 7T.

**Fig 2:**
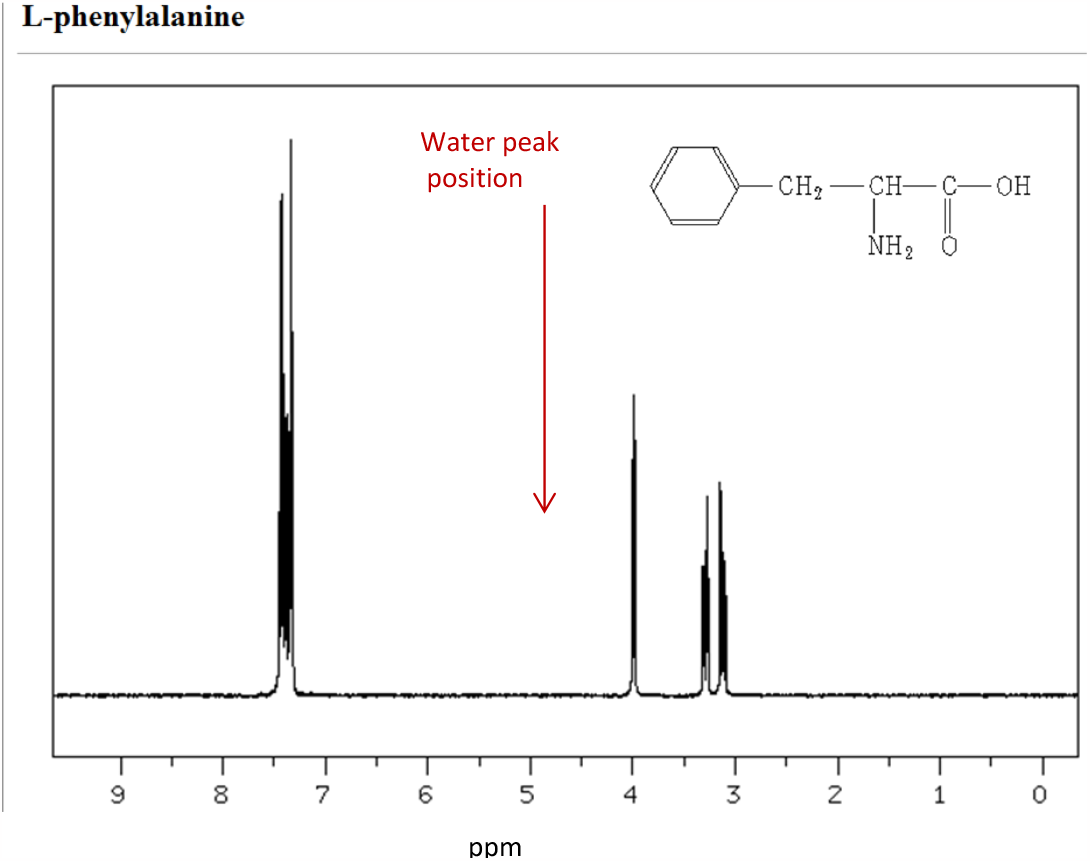
Spectrum of a phenylalanine solution in deuterated water (D_2_O). The water peak seen in vivo at 4.7 ppm is not present in this spectrum which is acquired from a D_2_O solution. The aim of the investigation is to detect the highest signal in the spectrum, which is to the left of the water peak position in vivo, at around 7.3 ppm. It arises from the phenyl ring of L-phenylalanine. Adapted from the spectral database for organic compounds SDBS: http://sdbs.db.aist.go.jp/sdbs/cgi-bin/direct_frame_top.cgi.

## MATERIALS and METHODS

### Phantom data Acquisition

In order to validate and optimize the MRS techniques used for observing phenylalanine in the brain at 7T, L-Phenylalanine solutions were prepared with 10 mM, 5 mM, 1.8 mM, 0.6 mM and 0.2 mM concentrations. The lowest concentration (0.2 mM) was selected based on the reported [4] phenylalanine concentration in the human brain. The phantom solutions were contained in 50 ml conical tubes. For data acquisition, each tube was immersed separately in a water bath and placed in the 32 channel Nova head coil. MRS measurements were obtained using single voxel spectroscopy (SVS) techniques optimized for 7T from a 40 x 15 x 15 mm^3^ voxel. We used STEAM (STimulated Echo Acquisition Mode) sequence with an ultra-short TE of 5 ms, TR=10 s and 32 averages, and the SVS sLASER (semi-adiabatic Localization by Adiabatic SElective Refocusing) sequence with a minimum TE of 26 ms.

### L-Phenylalanine measurements *in vivo* in humans

Having optimized phenylalanine signal observation in phantom solutions, we then attempted measurements in the human brain on 15 healthy volunteers and optimized the *in vivo* protocol parameters. We acquired downfield spectra in different areas of the brain with different voxel sizes. The absence of a distinct phenylalanine signal from normal brain concentration, led us to investigate variations in the metabolite spectral features upon increased level of brain phenylalanine, and to determine the concentration level needed for the phenylalanine signal to be detected *in vivo*. For practical reason this investigation was done on a cat brain as detailed below

### L-Phenylalanine measurements in cat brain

In order to determine the smallest phenylalanine concentration needed for *in vivo* detectability at 7T, spectra from cat brain were acquired following IV injection of phenylalanine. The cat weighed 4.12 Kg. with the cat positioned in the 32 channel Nova head coil, four Injections of 100mg/kg each were administered at one hour intervals. Prior to the IV injection, shimming and SNR were optimized on a (8x10x10) mm^3^ voxel. Because of its higher SNR, the SVS sLASER sequence was used with TR=4700 ms,TE=26 ms, 576 averages and a measurement time of 45 min. A base spectrum (Fig 5 left) was acquired before the first injection, and followed by four serial spectra acquired every hour right after the IV injection.

## RESULTS

### Phantom

The spectra in Fig. 3, obtained with the STEAM sequence, illustrate the phenylanaline multiplet at about 7.4 ppm. As expected, the lower concentration (0.2 mM) spectrum (left) has a much reduced SNR compared to the 10mM concentration spectrum (right). It is to be noted that for optimum detection of the downfield (4.7 ppm – 9 ppm) phenylalanine signal, the sequences were modified to allow shifting of the acquisition central frequency to 7.4 ppm in order to reduce loss of signal from the chemical shift displacement effect.

**Fig 3:**
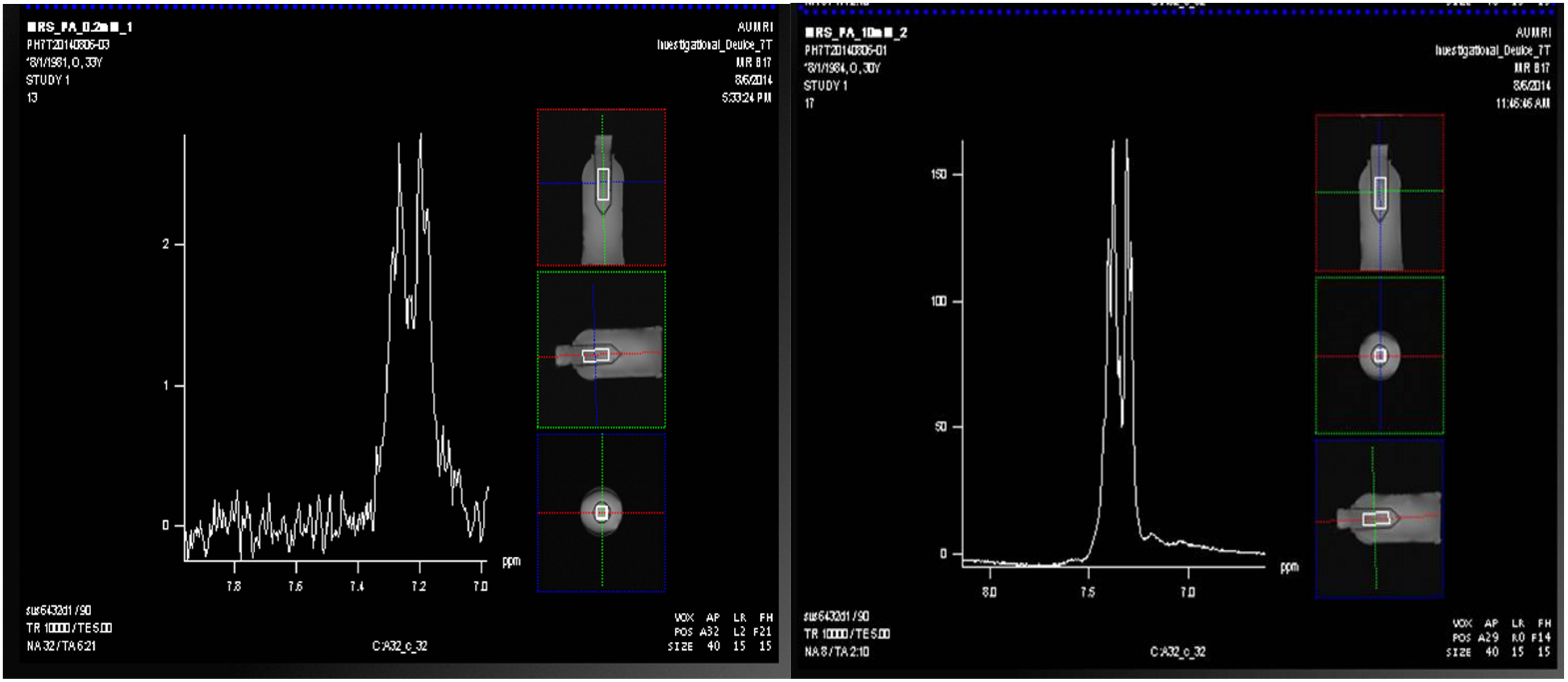
Phenylalanine peaks at 7.35 ppm from the 0.2 mM (left) and 10mM solutions (right).

### L-Phenylalanine *in vivo* in humans

Example spectra obtained *in vivo* from human volunteers are shown in Fig 4. with metabolite signals in the 6.5 – 9.00 ppm range. Those include the most dominant peak NAA at 7.8 ppm, a cluster of peaks between 6.5 and 7.5 ppm assigned to adenosine triphosphate (ATP,) glutamine (Gln), glutathione (GSH), phosphocreatine PCr as well as other moieties. If present at 7.3 ppm, Phe would overlap with PCr at 7.27 ppm and, DEPENDING on the shim, with other neighboring peaks. The absence of a distinct phenylalanine signal from normal brain concentration, led us to investigate variations in the metabolite spectral features upon increased level of brain phenylalanine, and to determine the concentration level needed for the phenylalanine signal to be detected *in vivo*. For practical reason this investigation was done on a cat brain as detailed below.

**Fig 4:**
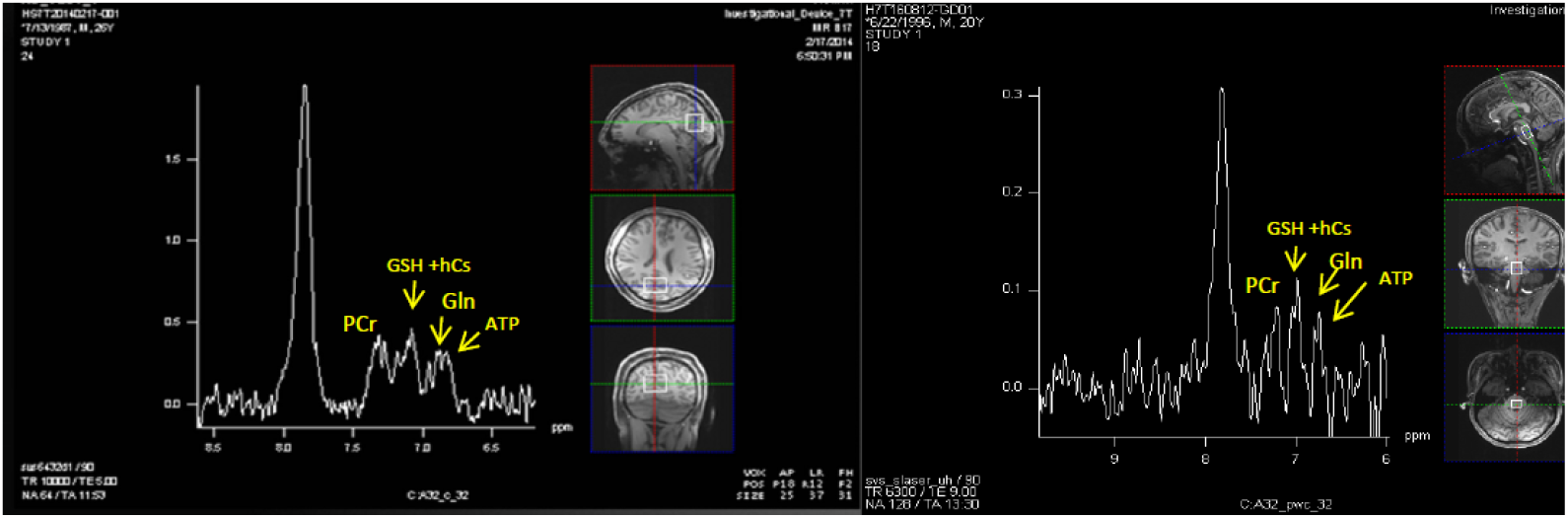
Examples of downfield spectra from the brain of healthy volunteers. The right spectrum is from a (11 x 16 x 22) mm^3^ voxel and has a much lower SNR compared to the left spectrum which was acquired from (25 x 37 x 31) mm^3^ voxel.

### L-Phenylalanine in cat brain

Fig 5 illustrates the base spectrum (left) and the final spectrum (to the right) after the 4 injections. Both spectra show the known NAA peak at 7.8 ppm. The remaining metabolites previously seen in the human brain are not as visible here due to the much smaller voxel ((8x10x10) mm^3^) used in the cat brain. As indicated by the arrows, no signal is detected at around 7.3 ppm in the base spectrum (Fig 5 left) and in the spectra following the 1^st^, 2^nd^ and 3^rd^ injections (not shown here). But a peak is seen at that position in the final spectrum following the 4 phenylalanine injections with a total of 400mg/Kg in the blood or a concentration of 3.4mM. Hence the wide peak at about 7.3 ppm may be assigned to the phenylalanine that has been accumulating in the brain over the 4 hour duration of the experiment. A similar experiment [4] performed in healthy volunteers showed that the brain phenylalanine concentration stays lower than blood phenylalanine concentration all through the phenylalanine oral loading process and approaches the blood concentration after 5 hours of loading at one hour intervals. Assuming a similar behavior in cat brain, this could explain the absence of the resonance peak at 7.3 ppm in the cat brain spectra following the 1^st^, 2^nd^ and 3^rd^ injections. After 4 hours of IV loading, the phenylalanine concentration in the cat brain may be close to the blood concentration of 3.4 mM. This would then imply that a phenylalanine concentration of 3.4 mM is needed for detection of phenylalanine from such a small voxel of brain tissue.

**Fig 5:**
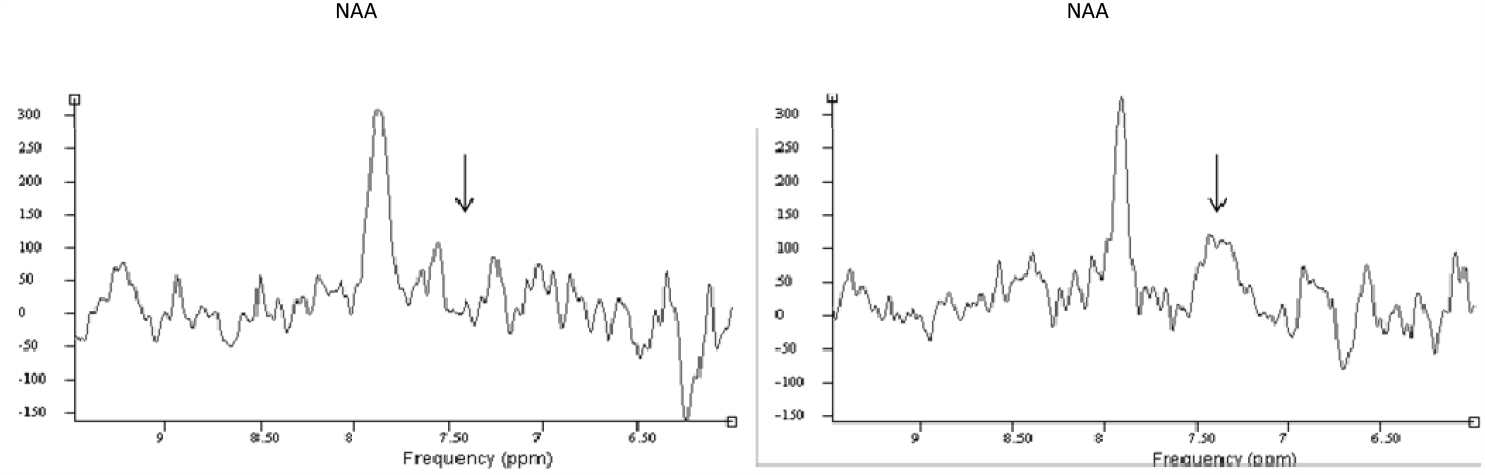
Spectra from a (8x10x10) mm^3^ voxel in a cat brain before (left) Phenylalanine IV injection and following four phenylalanine injections (right) of 100mg/kg each delivered 1 hour intervals. The arrows indicated the 7.3 ppm position where no signal is seen in the left spectrum and a wide peak assigned to phenylalanine is seen in the right spectrum.

## DISCUSSION and CONCLUSION

Phenylalanine concentration is about 0.2 mM in the normal brain and cannot be detected with MRS at lower field strength. Previous studies have measured elevated levels of phenylalanine in healthy volunteers [4] following oral phenylalanine loading and in the brain of patients with phenylketonuria [5], which is characterized by high levels of phenylalanine in the blood as well as in the brain. These studies were performed on lower field strength scanners (1.5T and 2T) with MRS spectra acquired from very large ROI (70 cm^3^). Our proposed study is an attempt at phenylalanine measurement in the brain using 1H MRS using an ultra-high field scanner (7T) and more technologically advanced multichannel MRI coils. It was expected that with the advantages of the 7T scanner (improved SNR and spectral resolution), we would be able to use smaller voxels to detect phenylalanine and distinguish it from neighboring overlapping peaks.

However our current investigation has confirmed that normal concentration of phenylalanine cannot be detected in the brain of healthy volunteers with MRS at 7T. These results are corroborated by a recent publication [6] which also concluded that normal phenylalanine is too small to contribute to the resonances around 7.3 ppm. We have detected phenylalanine from a very small voxel of the cat brain following 4 IV injections of 100mg/kg each; and we deduced that a brain tissue phenylalanine concentration of about 3.4 mM is needed in order to detect the phenylalanine signal from a 0.8 cm^3^ voxel. The motivation for using a small voxel is keeping with the expectation that one would want to measure phenylalanine in the locus coeruleus of the human brain, a small region with less than 2 cm^3^ volume, given its etiological role in Alzheimer’s disease. However, a phenylalanine concentration level 3.4 mM is not expected to be present in the brain of AD patients. With such a small voxel and the small phenylalanine concentration, it is to be concluded that phenylalanine cannot be unambiguously measured in the brain of AD patients or healthy controls with the currently available techniques on the 7T MRI scanner.

## References

1. P Herregodts, G Ebinger, and Y Michotte, “Distributions of monoamines in human brain: evidence for neurochemical heterogeneity in subcortical as well as cortical areas,” Brain Research, vol. 542, pp. 300–306, 1991.

2. D Weinshenker, “Functional Consequences of Locus Coeruleus Degeneration in Alzheimer’s disease,” Current Alzheimer Research, vol. 5, pp. 342–345, 2008.

3. MT Haneka et al., “Noradrenergic depletion increases inflammatory responses in brain: effects on IkappaB and HSP70 expression,” Journal of Neurochemistry, vol. 85, pp. 378–398, 2003

4. J Pietz, T Lutz, K Zwygart, GF Hoffmann, F Ebinger, C Boesch and R Kreis, “Phenylalanine can be detected in brain tissue of healthy subjects by 1H magnetic resonance spectroscopy”, J.Inherit.Metab. Dis 26, 683–691, 2003.

5. R Kreis, J Pietz, J Penzien, N Herschkowitz and C Boesch, “Identification of phenylalanine in the brain of patients with phenylketonuria by means of localized in vivo 1H Magnetic Resonance Spectroscopy”, Journal of Magnetic Resonance, series B, 242–251, 1995.

6. ND Fichtner, A Henning, N Zoelch, C Boesch and R Kreis, “Elucidation of the Downfield Spectrum of Human Brain at 7T Using Multiple inversion Recovery Delays and Echo Times”, Magnetic Resoance in Medicine, 78, 11–19, 2017.

